# Demographic variation and demographic niches of trees species in the Barro Colorado Forest

**DOI:** 10.1101/2022.07.07.499151

**Authors:** Richard Condit, Nadja Rüger

## Abstract

One goal for the 50-ha plot on Barro Colorado Island since its inception has been to understand demographic variability across the entire community of tree species. Early papers classified demographic response of many species to canopy gaps, culminating over the last decade with improved statistical methods that could quantify the response of growth, survival, and recruitment rates to increasing light for the entire community. We compile results from recent studies to illustrate the demographic trade-off between fast growth in high light and long life in deep shade. Growth and recruitment in high light are significantly correlated with mortality across species, and this trade-off is the primary demographic differentiation in the community. Here we add a new analysis of expected adult lifespan of 31 dominant canopy species spanning the growth-mortality axis. Using analytical solutions to matrices describing the lifetable, we demonstrate that the trade-off between high growth and long lifespan is not equalizing. The expected adult lifespan of a newly-recruited sapling is shorter in fast-growing pioneers than in long-lived shade-tolerant species: elevated growth rates of pioneers is insufficient to overcome their high death rates. If reproductive output were equal across the demographic axis, pioneer species could not persist. This suggests pioneers out-reproduce shade-tolerant species. The next goal is incorporating seed and seedling production into the demographic analysis to test the trade-off using lifetables having reproductive output included.

## Introduction

From the outset, the 50-ha plot on Barro Colorado Island included the goal of understanding how tree species demography responds to variation in light environment in the forest, thus testing how species partition the regeneration niche created by high light below canopy openings (Grubb 1977, Hartshorn 1980, Denslow 1980). A familiar paradigm in tree life history contrasts species that are tolerant of deep forest shade with those that regenerate only in clearings formed when tree dies. This contrast in life history offers a way for species to coexist along the light gradient from shade to small gap and large gap (Denslow 1980, Brokaw 1985, Pacala & Rees 1998). With a 50-ha census of more than 200,000 individual trees, Hubbell and Foster (1985) intended to assess life history niches in most of the forest species. In contrast, most attempts to examine life history variation in diverse tropical forests include only a handful of species (eg Brokaw 1987, Denslow 1990, Brown & Whitmore 1992, Kobe 1999, Svenning 2000, Dalling et al. 2004, Balderrama & Chazdon 2005).

Understanding the regeneration niche required detailed measurements of the light environment, and Hubbell and Foster (1986) included a map of light gaps from the outset of the project, estimating the density of vegetation above each corner of a 5×5 m grid over all 50 ha. Using this map, Welden et al. (1991) compared recruitment, growth, and survival rates in gaps versus shade for more than half the species in the forest. The definition of a gap in Welden et al. was strictly dichotomous: gaps were sites with no vegetation more than 10 m above the ground, and everywhere else was non-gap. In Condit et al. (1996), Welden’s index of gap preference based on recruitment was combined with growth and survival rates to assign tree species into demographic guilds reflecting shade-tolerance and growth rate, again including more than half the plot’s species.

Starting with Condit et al. (2006), we incorporated Bayesian methods to improve species-level estimates of demographic rates and to accurately estimate the variance of rates across the community. The use of Bayesian hyper-distributions allows rare species to be included, and since sampling error is modeled explicitly, extreme estimates in rare species are avoided. This leads to unbiased estimates of community-wide variation. We subsequently refined a three-dimensional map of light across the 50 hectares by integrating the vegetation profile at any point both horizontally and vertically, leading to a continuous estimate of vegetation density and thus light as a percent of full sun at every location (Rüger et al. 2009). Using the new map of light, Rüger et al. (2009, 2011ab, 2012) applied the Bayesian approach to describe recruitment, growth, and mortality as a continuous function of light for 90% of the species in the 50-ha plot. This allowed us to define niche axes based on complete demographic profiles (Rüger et al. 2018).

Here we summarize demographic axes that combine growth, mortality and recruitment, and we offer for the first time as a data supplement complete tables of the demographic parameters that describe species responses to light. These offer users demographic traits for close to 90% of the 300 species in the plot (Rüger et al. 2022). Many future studies of morphology, physiology, or population biology of Barro Colorado trees can start with these (Rüger et al. 2020).

We then evaluate the fast-slow continuum of Barro Colorado trees by asking whether it works as a demographic equalizing axis. This would be the case if high growth of pioneers overcomes their high death rate, so that saplings of pioneer and shade-tolerant species are equally likely to spend time at reproductive size. If so, and given equal reproductive output, the two groups of species would have equal population-level fitness. It appears to us that the notion of a fast-slow demographic axis (Salguero-Gomez 2017) includes the equalizing assumption implicitly. Here we offer a rigorous test of the assumption by estimating the reproductive lifetime of the average sapling in 31 abundant species, integrating the growth and survival schedules using a population matrix.

### Demographic responses to light

We start by combining the results from Rüger et al. (2009, 2011ab) into axes of mortality-recruitment and mortality-growth (Fig. 1). The figures make it clear that there are no separate demographic categories: pioneers are simply species at the upper end of continuous variation, including the abundant *Miconia argentea* and *Cordia bicolor* (Fig. 1). Conversely, shade-tolerant species, such as *Prioria copaifera* and *Swartzia simplex*, are the low end. Between those extremes, species are spread along the axis in ways that had not been appreciated. Forest dominants such as *Trichilia tuberculata, Tabernaemontana arborea, Beilschmiedia tovarensis, Guatteria lucens*, and *Guarea guidonia*, all recruiting abundantly in the forest understory, are distinct in life history (Fig. 1).

**Figure 1.**
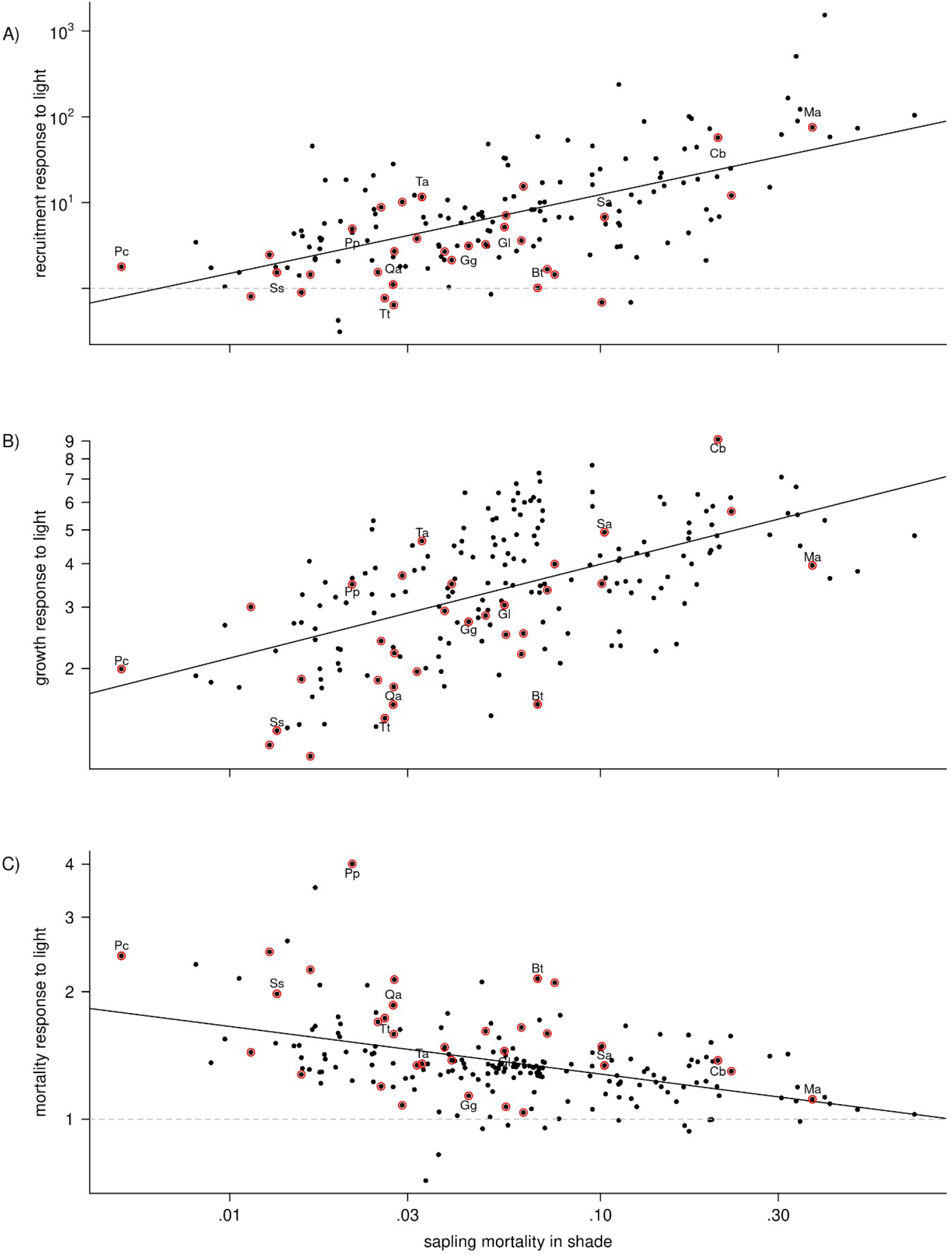
A) Recruitment response to light versus mortality rate in the shade for 165 tree species (taller than 12 m) from the Barro Colorado 50-ha plot. B) Growth response of 1-cm saplings to light versus mortality rate in the shade for 197 species. C) Mortality response of 1-cm saplings to light versus mortality rate in the shade for 203 species in the plot; 163 species appear on all 3 panels. The vertical axes are ratios of recruit density, dbh growth, or mortality at 20% versus 2% light, so values > 1 mean higher rates in high light. Mortality is the death rate of 1-cm saplings at 2% light. There is a horizontal dashed line at 1, indicating no response (it is outside the graph in B). In A) and C), a few species have responses < 1, indicating higher recruitment (mortality) in the shade, but no species had higher growth in the shade. Axes are logarithmic. Solid lines are regressions, and all three were highly significant (based on log-transformed values). The 31 species used in lifetable calculations are highlighted by red circles, and 11 are identified: Cd *= Cordia bicolor*, Gg = *Guarea guidonia*, Gl *= Guatteria lucens* (formerly *G. dumetorum*), Ma = *Miconia argentea*, Pc = *Prioria copaifera*, Pp = *Protium panamense*, Qa = *Quararibea asterolepis*, Sa = *Simarouba amara*, Ss *= Swartzia simplex var. grandifolia*, Ta = *Tabernaemontana arborea*, and Tt *= Trichilia tuberculata*.

At the pioneer end of the axis, there are species having recruitment rates increasing > 100 fold from shade (2% light) to high light (20%), and growth rates increasing by > 6-fold (Fig. 1). But both recruitment and growth increased with light in nearly every species in the forest (Fig. 1). Even the tree species we know as shade-tolerant, including abundant canopy dominants, had recruitment and growth rates 2-4 fold higher in high light relative to shade (Fig. 1). We included a map in the supplement to Rüger et al. (2009) showing recruits of all species overlain on a map of high light areas, and forest-wide recruitment is conspicuously concentrated in high light.

Previous studies on mortality of tropical trees in gap versus non-gap often reported higher mortality in shade (Augspurger 1984, Kobe 1999, Balderrama & Chazdon 2005), but some results suggested the opposite (Fraver et al. 1998); all considered seedlings only. The early study on gaps in the Barro Colorado plot (Welden et al. 1991) suggested higher mortality in gaps, though individual species did not have statistically significant responses. Our new results support Welden et al. (1991), demonstrating increased mortality with light in most species (Rüger et al. 2011). The mortality response to light was negatively correlated with growth and recruitment responses, meaning species with strong growth and recruitment responses tended to have smaller mortality responses, but mortality was elevated in response to light even in those light-demanding pioneers (Fig. 1C). A few highly shade-tolerant species, such as *Protium panamensis* and *Prioria copaifera*, had mortality rates > 2-fold higher in 20% light compared to shade. Elevated mortality in high light could be caused by damage from tree or branch falls in gaps, and pioneer species might avoid the effect because they invade high light sites and grow upward quickly, avoiding the worst of the branch falls.

The correlation lines in Figure 1 depict primary axes of demographic variation, from slow to fast species (Stearns 1999, Salguero-Gómez 2017), a trade-off between survival in the shade and growth (or recruitment) in high light (Reich 2014, Rüger et al. 2018, 2020). There is a physiological understanding of the trade-off, since the shift to high growth requires traits that mitigate against high survival (Kitajima 1994, Poorter 1999, Sack & Grubb 2001, Wright et al. 2010, Philipson et al. 2014). Physiology alone, however, cannot, explain community dynamics, and what the fast-slow axis means to population fitness, or net demographic performance, is crucial. An analysis of lifetables constructed from observed growth and mortality will account for continuous variation along the axis as well as the spread above and below the axis.

### Variance around the main axis

Rüger et al. (2018, 2020) extended observations of demographic trade-offs by considering variation around the primary axis of fast vs. slow demography. The Bayesian analysis accounts for the variance caused by sampling error, which is high in rare species, and the remaining variance that Figure 1 reveals is the true difference between species. Despite the importance of the growth-mortality trade-off, many species fall far off the line, such as *Alseis, Protium*, and *Tabernaemontana* above versus *Beilschmiedia, Quararibea*, and *Trichilia* below (labeled in Fig. 1B). In terms of net demographic performance, those two groups are expected to be very different.

### Integrating demographic variation

On the surface, the qualitative link between high growth and high death might appear to equalize the net fitness of all the species, where fitness refers to integrated lifetime per capita population growth. But fitness is a quantitative trait requiring quantitative comparisons of lifetime demography across the entire spectrum of variation (Fig. 1). Here we preview a matrix population analysis that provides a test based on demography of trees above 1 cm dbh. A detailed exposition and tests for robustness of the conclusions are given in Condit (2022).

A matrix approach to demography is an analysis of a Markov transition process that allows the size distribution of trees in a population to be projected through time, allowing the maturation time from sapling to adulthood and the length of the reproductive period to be quantified (Caswell 2001). It requires describing the state of the population at one time as the number of individuals in several dbh categories, and a transition matrix that converts the population state from one time to the next. Define the population of trees in size class *i* as *N*_*i*_, the vector of *N*_*i*_ as **N**, and the transition matrix

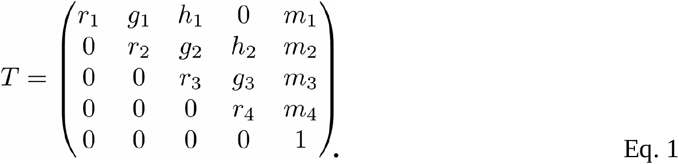

is defined so that **N’**=**N***T*. That is, multiplying current population **N** by *T* produces the population vector one time step later, **N’**. The matrix of Equation 1 covers four size classes, plus a fifth class for dead trees, ie *N*_*5*_ means the number dead. The matrix entries can be calculated from observed stem growth in repeat censuses: *r*_*i*_ terms are probabilities that a tree currently in size class *i* remains in size class *i* at the next census, *g*_*i*_ are probabilities that a tree in class *i* grows to class *i+1, h*_*i*_ means growth by two classes, and *m*_*i*_ are probabilities that a tree in class *i* dies in the next step. Class 4, the adult class, is terminal, with trees either remaining there (*r*_*4*_) or dying (*m*_*4*_). There is a 1 in the lower right because a tree in class 5, the dead class, has probability 1 of remaining dead. As an example, here is a 4-class transition matrix for *Trichilia tuberculata*:

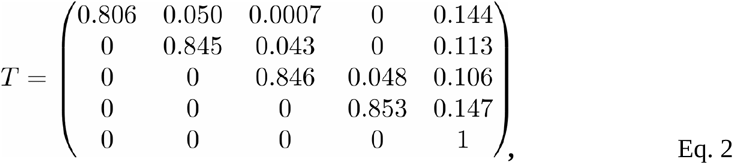

where probabilities mean 5-year transitions. The diameter classes are 10-34.4, 34.4-76.8, 76.8-200, and 200+ mm; 200 mm was set as the approximate reproductive size in *Trichilia* (Visser et al. 2016), and the other sizes were chosen by an algorithm designed to equalize the growth probabilities *g*_*i*_ as closely as possible. The estimates were calculated from observed stem transitions in each five-year census interval of the 50-ha plot from 1985-2015, then averaged across the six intervals. Entries below the diagonal, those transitions representing a tree moving backward to a smaller class, are zero, though in fact there were a few of cases where a small dbh error led to a tree shrinking below a class boundary. All those were assumed instead to be trees that remained in their earlier size class.

The advantage of the matrix approach is that it offers a direct calculation of one crucial demographic statistic: the expected adult lifespan of the average sapling. In the 4-class model, it is the expected number of time steps a sapling starting in class 1 will spend as an adult, in class 4. This average must include the many saplings that die before adulthood, so it must include the average adulthood of those that do make it discounted by zeroes for all those that do not. The time as a reproductive adult is crucial because, if the population is to persist, the length of time a sapling will spend reproducing must be sufficient to replace itself. If the primary axis of demographic variation, from slow to fast species (Fig. 1B), is also an axis of demographic equalization, then the expected adult lifespan should be equal across the axis.

The expected adult lifespan has a straightforward analytic formulation in the absence of two-step growth (*h=*0) and assuming no shrinkage. Define *L*_*j*_ as the expected lifespan in the terminal size class of a sapling starting in class 1. Then based on a matrix like Equation 1, but with an arbitrary number of size classes *j* (ie, *j* can be > 4),

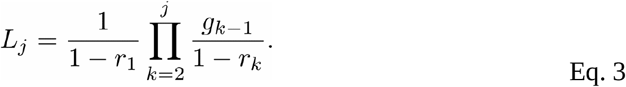

There is an analytical solution that includes *h* as well, though it does not generalize easily beyond *j=*5. Another useful result is the probability that a sapling reaches adulthood, and it has an analytical formulation,

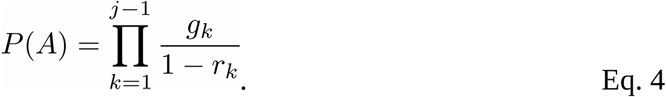

A third result is the maturation time, *M*, or the mean number of steps required for a sapling to reach adulthood. There is not an analytical formulation for *M* from the matrix in Equation 1, but there is for a simplified matrix, in which *g*_*i*_ and *r*_*i*_ are constant. Then, with *r*_*i*_ *=r*,

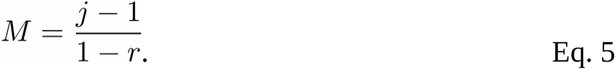

*L*_*j*_, *P(A)*, and *M* can also be calculated numerically by raising the transition matrix, *T*, to successive powers, since *T*^*X*^ is the transition matrix for *x* time steps (Kemeny & Snell 1976, Condit 2022). Complete derivations of Equations 3-5 are given in Condit (2022).

For the *Trichilia* transition matrix (Eq. 2), *L*_*4*_=0.15 time steps, which is 0.75 years since there are 5 years per census interval. This means that the average *Trichilia* sapling can expect to live less than 1 year as an adult. Analytical solutions for the probability of reaching adulthood is *P(A)=*0.022, and maturation time *M=*18.1 time steps (90.5 y), the latter found by setting *r* to the mean of the diagonal, *r*_*i*_ (Eqs. 1,2). Numerical solutions confirm the analytical results closely.

I calculated the transition matrix for the 31 most abundant species in the 50-ha plot that reach 16 m in height. All had populations of > 750 trees below 100 mm dbh and > 80 trees 100 mm or above in 1990, and had at least one tree taller than 16 m (Wright et al. 2010). Those cutoffs were chosen to include four fast-growing species, ensuring that the 31 species are spread widely across the growth-mortality axis (Fig. 1B). For those species with reproductive size of 300 mm dbh, I used 5 size classes instead of 4, always choosing the breaks between size classes to equalize growth transitions as closely as possible. In two fast-growing species, *Miconia argentea* and *Cordia bicolor*, there were two-step growth transitions fairly frequently, and for those I used the two-step version of Equation 3; in every other species, two-step transition probabilities were always < 0.05 of one-step probabilities. I estimated maturation time *M* using the numerical method, which has no limitations about two-step transitions.

### Adult lifespan and the growth-survival trade-off

The fast-growing pioneers performed poorly in the integrated demographic assessment, since they had short adult lifespans (Fig. 2A). A sapling of *Miconia argentea*, at the extreme, can expect to survive 6 months as an adult, and *Simarouba* and *Inga* < 2 years (Table 1). *Cordia*, on the other hand, had nearly 6 years expected adult lifetime (Table 1), demonstrating that pioneers can overcome high mortality with very fast growth (Fig. 1B). Several of the species at the slow end of the axis, *Prioria, Swartzia*, and *Heisteria concinna*, had expected adult lifespans > 20 years. For many of the commonest Barro Colorado tree species, those in the center of the growth-mortality axis, expected adult lifespan was highly variable. Some, especially *Trichilia* and *Guatteria*, had expected adult lifespans < 1 year, as low as pioneers, while *Alseis, Quararibea* and *Tabernaemontana* had lifespans > 10 years.

**Table 1.** Demographic results for 31 dominant species of the Barro Colorado 50-ha plot. Adult size is estimated reproductive size in mm dbh. *N* is the total number of individuals in the 1990 census. The x-axis is the position along the growth-mortality regression (Fig. 1), with high values for the fastest growing species. Adult expectations: *Lifespan* is the expected adult lifespan for the average sapling of 1 cm dbh (Fig. 2A); *Probability* is the fraction of those saplings that live to adulthood (Fig. 2B); *Maturation* is the mean time it takes a successful sapling to reach adulthood (Fig. 2C). *Swartzia simplex* includes one of the two varieties distinguished in the plot, *S. s. var. grandifolia*. The four pioneer species deliberately included by judicious choice of the abundance criterion were *Cordia bicolor, Inga marginata, Miconia argentea*, and *Simarouba amara*.

| Species (Family) | N | Adult size | Demographic axis | Adult expectations from 1 cm |  |  |
| --- | --- | --- | --- | --- | --- | --- |
|  |  |  |  | Lifespan | Probability | Maturation |
| *Alseis blackiana* (Rubiaceae) | 8412 | 100 | -0.14 | 10.74 | 0.072 | 124.9 |
| *Beilschmiedia towarensis* (Lauraceae) | 2748 | 100 | 0.32 | 1.35 | 0.023 | 117.4 |
| *Brosimum alicastrum* (Moraceae) | 920 | 100 | -0.48 | 4.85 | 0.080 | 188.3 |
| *Chrysophyllum argenteum* (Sapotaceae) | 688 | 100 | -0.21 | 3.03 | 0.025 | 137.3 |
| *Cordia bicolor* (Cordiaceae) | 1043 | 100 | 0.92 | 5.57 | 0.099 | 24.3 |
| *Cordia lasiocalyx* (Cordiaceae) | 1695 | 100 | 0.26 | 1.84 | 0.078 | 48.1 |
| *Drypetes standleyi* (Putranjivaceae) | 2279 | 100 | -0.17 | 15.53 | 0.140 | 107.0 |
| *Eugenia coloradoensis* (Myrtaceae) | 838 | 100 | -0.09 | 1.21 | 0.051 | 74.5 |
| *Eugenia oerstediana* (Myrtaceae) | 2343 | 100 | 0.00 | 2.28 | 0.085 | 52.4 |
| *Garcinia recondita* (Clusiaceae) | 4289 | 100 | -0.45 | 8.31 | 0.181 | 99.9 |
| *Guarea guidonia* (Meliaceae) | 1963 | 100 | 0.08 | 9.91 | 0.146 | 67.9 |
| *Guatteria lucens* (Annonaceae) | 1469 | 150 | 0.20 | 0.64 | 0.022 | 56.3 |
| *Heisteria concinna* (Olacaceae) | 984 | 200 | -0.58 | 50.49 | 0.548 | 61.8 |
| *Hirtella triandra* (Chrysobalanaceae) | 5026 | 200 | -0.22 | 22.58 | 0.282 | 70.6 |
| *Inga marginata* (Fabaceae) | 717 | 200 | 0.96 | 1.14 | 0.066 | 27.9 |
| *Lonchocarpus heptaphyllus* (Fabaceae) | 894 | 200 | 0.27 | 0.52 | 0.022 | 94.9 |

Table 1 (cont.).
| Species (Family) | N | Adult<br>size | Demographic<br>axis | Adult expectations from 1 cm |  |  |
| --- | --- | --- | --- | --- | --- | --- |
|  |  |  |  | Lifespan | Probability | Maturation |
| *Maquira guianensis* (Moraceae) | 1502 | 200 | -0.64 | 9.87 | 0.298 | 78.7 |
| *Miconia argentea* (Melastomataceae) | 900 | 200 | 1.24 | 0.54 | 0.033 | 18.8 |
| *Ocotea whitei* (Lauraceae) | 754 | 200 | 0.53 | 1.66 | 0.034 | 53.0 |
| *Poulsenia armata* (Moraceae) | 2120 | 200 | 0.37 | 0.34 | 0.014 | 60.6 |
| *Pouteria reticulata* (Sapotaceae) | 1767 | 300 | 0.21 | 5.28 | 0.044 | 73.7 |
| *Prioria copaifera* (Fabaceae) | 1439 | 300 | -1.08 | 31.52 | 0.219 | 151.1 |
| *Protium tenuifolium* (Burseraceae) | 3015 | 300 | 0.03 | 11.01 | 0.207 | 58.6 |
| *Quararibea asterolepis* (Malvaceae) | 2349 | 300 | -0.17 | 13.38 | 0.171 | 118.5 |
| *Simarouba amara* (Simaroubaceae) | 1289 | 400 | 0.54 | 1.28 | 0.050 | 37.6 |
| *Swartzia simplex* (Fabaceae) | 2573 | 100 | -0.56 | 42.79 | 0.301 | 138.1 |
| *Tabernaemontana arborea* (Apocynaceae) | 1430 | 300 | -0.07 | 14.73 | 0.255 | 69.7 |
| *Protium stevensonii* (Burseraceae) | 4114 | 300 | -0.16 | 5.03 | 0.057 | 109.2 |
| *Trichilia tuberculata* (Meliaceae) | 13284 | 300 | -0.20 | 0.75 | 0.022 | 90.3 |
| *Unonopsis pittieri* (Annonaceae) | 790 | 100 | 0.35 | 6.74 | 0.154 | 47.3 |
| *Virola sebifera* (Myristicaceae) | 2080 | 200 | 0.14 | 8.12 | 0.210 | 47.6 |

**Figure 2.**
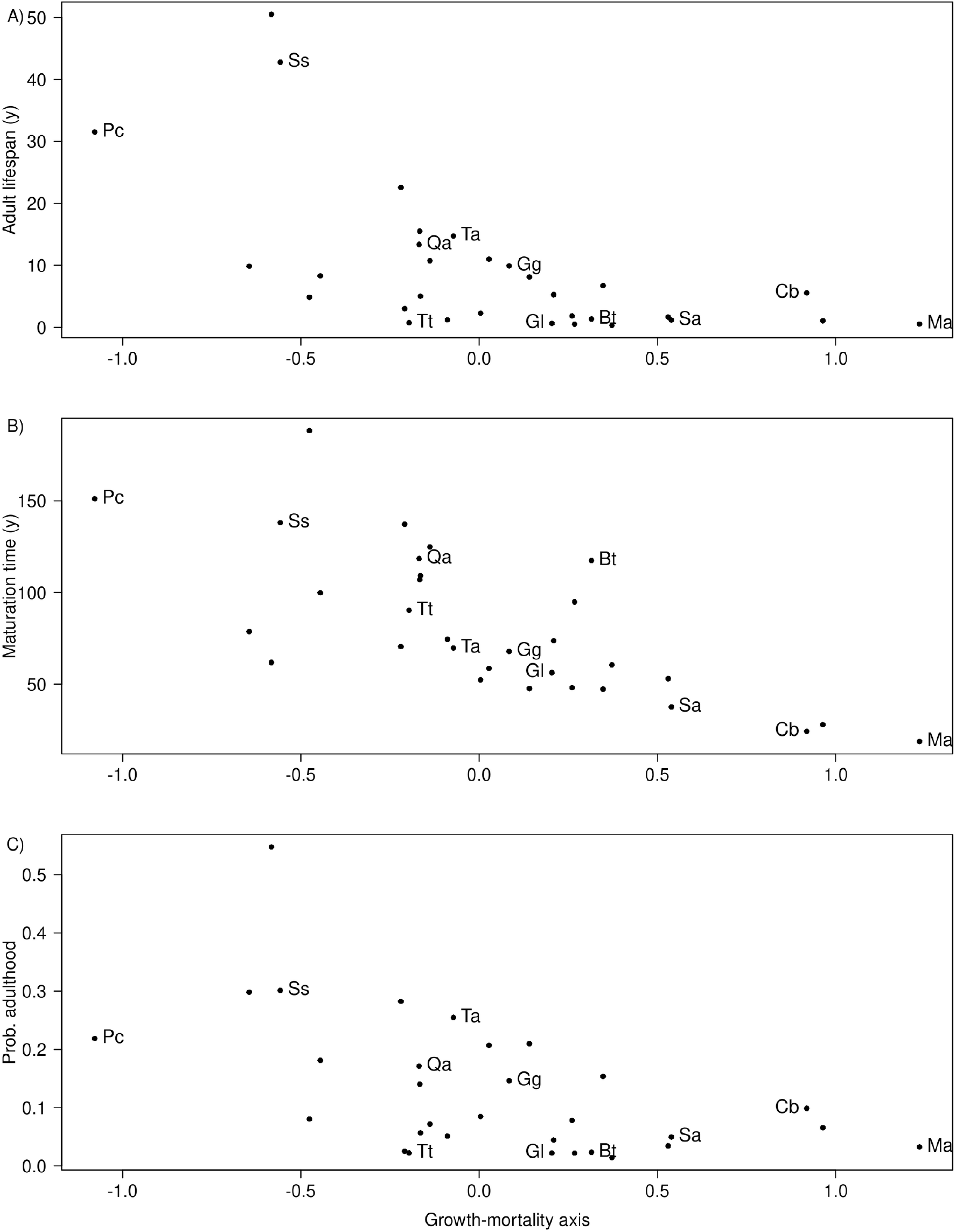
Lifetime demographic results (vertical axes) as a function of species’ positions on the growth-mortality axis (horizontal axes) for 31 abundant tree species chosen to represent the full range of the axis. The position along the axis was calculated from the regression line (Fig. 1B): first the predicted growth response was calculated for each species from the regression, then the distance of that prediction from the overall mean growth response was calculated as the species’ position. High positions are the fastest pioneers, to the right in Figure 1B. A) Vertical axis is the expected adult lifespan (years) for a sapling calculated from the transition matrix (Eq. 2); in two fast-growing species, the two-growth version of the equation was used. B) Vertical axis is the probability a sapling reaches adulthood. C) Vertical axis is the mean time (years) it takes for a successful sapling to reach adulthood. The 11 species identified are the same as in Figure 1.

Pioneers matured rapidly, and expected time for a sapling to reach adult size was < 20 years in *Miconia*, contrasting with *Prioria* and *Swartzia*, which needed > 140 years to reach adulthood (Table 1, Fig. 2B). The probability of reaching adulthood was low in pioneers and high in the long-lived species, but species in the middle of the axis were highly variable, and *Trichilia* had a lower probability of reaching adulthood than several pioneers (Fig. 2C).

## Conclusions

The demographic axis describes a growth-mortality trade-off that is not demographically equalizing. The expected adult lifespan of saplings in species at the fast end of the axis, the pioneers, is short relative to many slow-growing species. Though pioneers grow rapidly to adulthood, it is not quickly enough to overcome high mortality rates. In order to persist in the forest, the pioneers must have a reproductive advantage, ie by producing more seedlings and saplings during their reproductive life than long-lived species do. Rüger et al. (2018, 2020) and Kambach et al. (2022) pursued this possibility by including recruitment rate at the 1 cm size in analyses of demographic axes. This leads to the suggestion that there are species below the growth-mortality axis of Figure 1 that persist in the forest by recruiting well (Rüger et al. 2018). Alternatively, it may be that the pioneer species are so concentrated in areas of high light that the only demography that matters is their maximum growth, not the average growth across the plot. Perhaps a lifetable constructed from only those saplings in high light would lead to sufficient adult lifetimes. Moreover, we know that the community of species abundances in the 50-ha plot is far from stable, with many species increasing or decreasing by substantial amounts (Chisholm et al. 2014, Condit et al. 2017), and the variation in adult lifespan revealed in Figure 2 should account for those fluctuations.

An overarching question is whether the demographic trade-off axis is stabilizing in the sense that species abundances are held in a narrow range, thus fostering coexistence (Grubb 1977, Hubbell et al. 1999, Chesson 2000, Silverton 2004). Stable coexistence of two species, one at the upper end of the axis and one at the lower, is easy to understand, and demographic variation may be an important coexistence mechanism in low-diversity temperate forests. The question at Barro Colorado is whether coexistence of hundreds of species based on a demographic trade-off is plausible. Tilman (2004) argued that many species spread uniformly along a single resource axis could persist in a stable community. The principal argument against this hypothesis is that stabilizing differences among hundreds of species would be trivially small, almost certainly smaller than population drift (Condit et al. 2012). Moreover, the observed spacing of species along the demographic axis counters the Tilman prediction, since most species are clustered in the center (Fig. 1; Condit et al. 2006). Even with a second axis of moisture variation within the Barro Colorado plot, there are too many species for stable coexistence via local niche partitioning. At much wider scales, the issue reopens, because climatic and soil variation are much greater, and species disperse widely. Species that are demographically equivalent within the Barro Colorado plot could be maintained by dispersal from sites where they differ (Condit et al. 2013).

Future studies need to go beyond the documentation of species differences and focus on demographic consequences of those differences. Our analysis here moves in that direction, and the further step of incorporating reproduction and seedling demography into the lifetables must follow. There are now 30 years of seed and seedling demography (Wright et al. 2005, 2015) to go with 30 years of the main census (Condit et al. 2017). Combining them can provide the precise estimates of lifetime demography needed for understanding the consequences of species differences. Meantime, we offer now, for the first time, complete demographic profiles calculated in Rüger et al. (2009, 2011ab) and thus thorough documentation of observed species variation (Condit et al. 2022).

